# Isolation of an infectious mammalian chu-like virus from tumor cells of the endangered Tasmanian devil (*Sarcophilus harrisii*)

**DOI:** 10.1101/2024.11.25.625296

**Authors:** Julien Mélade, Erin Harvey, Jackie E. Mahar, Jocelyn M. Darby, Andrew S. Flies, Edward C. Holmes

## Abstract

*Jingchuvirales* (negative-sense RNA viruses) were initially discovered in invertebrates, with both exogenous and endogenous jingchuviruses subsequently identified in fish, reptiles and mammals. To date, jingchuviruses have only been described metagenomically. By screening primary tumor tissues and tumor cell lines from the endangered Tasmanian devil (*Sarcophilus harrisii*), we isolated Tasmanian devil chu-like virus (TDCV) from cultures of Tasmanian devil facial tumor disease (DFTD) cells. Cell infection experiments demonstrated active virus replication in Tasmanian devil tumor cells, but not mosquito cells. The absence of viral replication in fibroblasts in cell culture and the lack of RNA detection in several organs suggested that replication was associated with tumor cells. Phylogenetic analysis revealed that TDCV likely represents a novel virus family. This is the first isolation of a jingchuvirus, demonstrating their capacity to infect mammalian cells, and providing *in vitro* avenues to understand the biology of TDCV and its association with tumor cell infection.

## INTRODUCTION

The order *Jingchuvirales* are a diverse group of single-stranded negative-sense RNA viruses currently classified into five families, including the *Chuviridae* from which their name was derived^1^ as well as currently unclassified “chu-like” viruses. The *Jingchuvirales* are characterized by remarkable genomic plasticity, comprising segmented, non-segmented, linear and circular genomes, which range in length from 9.1 to 15.3 kb^2^. The *Chuviridae* and *Aliusviridae* families within the order each contain several viral genera and species, while the *Natareviridae*, *Crepuscuviridae* and *Myriaviridae* currently possess only a single species each^1^.

Jingchuviruses were discovered through the metagenomic sequencing of arthropods and were initially believed to be exclusively associated with invertebrates^3^. More recently, however, jingchuviruses have been identified in both fish and reptiles^4–9^, and chu-like viruses have been sporadically detected in tissue samples from small mammals^10,11^ as well in as a basal chordate (a tunicate)^12^. Of note, the chu-like sequences detected in rodents and shrews in China^10^ were confirmed by PCR as present in multiple organs (gut, liver, lung spleen and kidney), with abundance values reaching 681 reads per million (RPM), suggestive of active replication in mammalian cells. In addition, endogenous viral elements (EVEs) of jingchuviruses have been identified in the genomes of ray-finned fish^4^, as well as both placental mammals and marsupials^13^.

The chuviruses identified in vertebrates through metagenomic studies of fish and reptiles form a monophyletic group and are classified within a single genus - *Piscichuvirus* - within the *Chuviridae*. However, phylogenetic analysis revealed that the endogenous and exogenous viral sequences from fish, reptiles, amphibians and marsupials were topologically interspersed within the genus and did not form distinct groups. Hence, there have likely been multiple virus endogenization events in vertebrates over an unknown evolutionary time-scale^14^. Similarly, the RNA-dependent RNA polymerase (RdRp) protein sequences from the chu-like viruses found in small mammals did not form a monophyletic group and did not cluster with the piscichuviruses^10^. Instead, they fell in diverse locations throughout the *Jingchuvirales* phylogeny^10^, indicating that there have been several independent emergence events in mammals, likely from invertebrate hosts. A jingchuvirus EVE recently identified in the bottlenose dolphin (*Tursiops truncatus*) was also phylogenetically distinct from the piscichuviruses, reflecting another cross-species transmission event^13^. Hence, jingchuviruses have more complex evolutionary histories and host associations than deduced from their initial detection in arthropods^3,4^. To date, however, no jingchuvirus has been cultured or isolated, such that very little is known about the underlying biology or true hosts of this rapidly expanding viral order.

By analyzing all RNA-sequencing (i.e., metatranscriptomic) data publicly available in the NCBI Sequence Read Archive (SRA) for the Dasyurid order of marsupials we recently identified Tasmanian devil chu-like virus (TDCV) in a Tasmanian devil (*Sarcophilus harrisii*) tumor cell line^11^. TDCV has a non-segmented, non-circular genome, and its high abundance (0.2% of total reads) suggested a strong viral replication in these cells. TDCV is of particular interest as the Tasmanian devil is an iconic and endangered species only found on the island of Tasmania, and over the last 20 years has experienced regional population declines averaging 82%^15^. The main reason for this dramatic population reduction is the spread of two independent transmissible cancerous clones - devil facial tumor 1 (DFT1)^16^ and devil facial tumor 2 (DFT2)^17^. Both cancers usually manifest on the face and oral cavity, and lead to death via metastatic disease or starvation. Schwann cells of the peripheral nervous system are the origin of the tumoral cells in this disease^18^. Importantly, TDCV was metagenomically detected in a cell line established from DFT1 cells. However, the true host association and the ability of this virus to replicate in these cells remains to be confirmed.

Herein, we screened for the presence of jingchuvirus RNA both in primary tissues from individuals with Tasmanian Devil Facial Tumor Disease (DFTD) and cell lines established from tumor tissues. Using cell culture experiments we examined the replicative capacity of TDCV in both marsupial tumor cell lines and arthropod cell lines, and performed phylogenetic analyses to reveal the evolutionary origins of this virus and the vertebrate jingchuviruses as a whole.

## RESULTS

### RT-PCR and RNA-seq screening for TDCV and chu-like virus RNA

A total of 98 primary tissues representing spleen (n = 28), liver (n = 18), kidney (n= 13), colon (n = 1), and lymph nodes (n = 1) collected from Tasmanian devils, as well as 37 Tasmanian devil primary facial tumoral tissues and 8 cell lines (tumoral and fibroblast), were screened for the presence of chu-like viruses using an end-point RT-PCR (**Supplementary Table 1 and 2**). Notably, one of the cell lines tested here, DFT1 4906, corresponds to the same cell line that was sequenced and uploaded to the SRA in which we first identified

TDCV in a previous study^11^. As expected, we observed PCR amplification of TDCV in DFT1 4906, which is a Tasmanian devil tumor cell line previously established from DFT1 facial tumor tissue (**Figure 1**). Based on qRT-PCR, the number of RNA copies in the sample was

**Figure 1.**
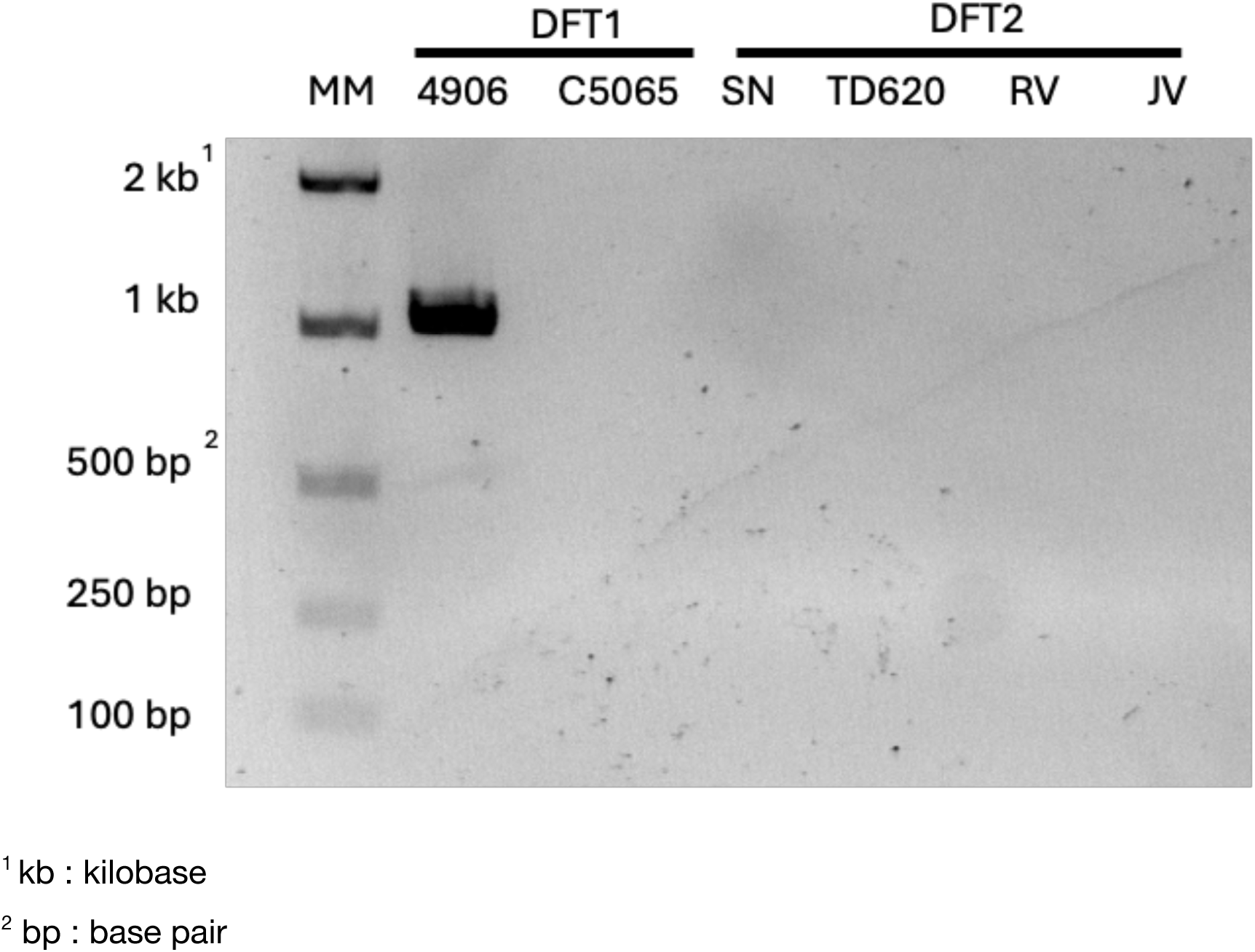
RNA amplification of Tasmanian Devil chu-like virus (TDCV) in the seven DFT cell lines by RT-PCR. RNA extracts from DFT1 and DFT2 cell lines were amplified by RT-PCR and amplicons were run on an electrophoresis gel. Labels at the top of the gel image indicate the cell line (M: Molecular weight ladder).

7.2 log10 copies/mL, + /- 0.1 log10 copies/mL. We did not observe amplification of TDCV in other cell lines, organ tissues or tumoral primary tissues. Unbiased RNA sequencing was also used on the same samples to identify other jinchuvirales: the spleen, kidney, liver, colon and lymph nodes in our Tasmanian devil samples were negative for any chu-like virus. Equally, no additional chu-like viruses were found in the cell lines, aside from in DFT1 4906, in which TDCV was found in high abundance as expected. However, RNA-seq analysis identified an abundance of viral sequences in each tumoral cell line corresponding to bovine viral diarrhea virus (BVDV), a frequent contaminant of fetal bovine serum used in cell cultures^19^.

### Detection of viral negative and positive strands

To determine if TDCV is capable of efficient replication in the tumor cell line, both the positive and negative strands of the virus, that act as markers of active replication, were amplified and quantified by RT-qPCR. From this, we observed concentrations of 6.5 and 7.0 log10 copies/mL (+ /- 0.05 log10 copies/mL) for the positive and negative strands, respectively, in the DFT1 4906 tumoral cell supernatant, indicative of replication.

### Characteristics of infected and non-infected tumoral cell lines

The six tumor cell lines, including infected DFT1 4906, were defrosted and cell viability was recorded at confluency for each passage. In the first passage, microscopic analysis revealed that the morphology of the cell line DFT1 4906 naturally infected by TDCV was markedly different from the other non-infected tumor cell lines, including a different DFT1 cell line, C5065 (**Figure 2**). While uninfected cells presented a fibroblastic morphology, infected DFT1 4906 cells were small with a round shape. To analyze the relative fitness of infected DFT1 4906 cell lines and uninfected cell lines, cells were passage no more than 15 times and cell proliferation was quantified daily. Interestingly, we noted a significant proliferation reduction in the infected DFT1 4906 cell density compared to the uninfected tumoral cell lines (**Figure 3**) after 13 passages, but observed no reduction in TDCV RNA copy number in the cell supernatant from naturally infected DFT1 4906 at each passage.

**Figure 2.**
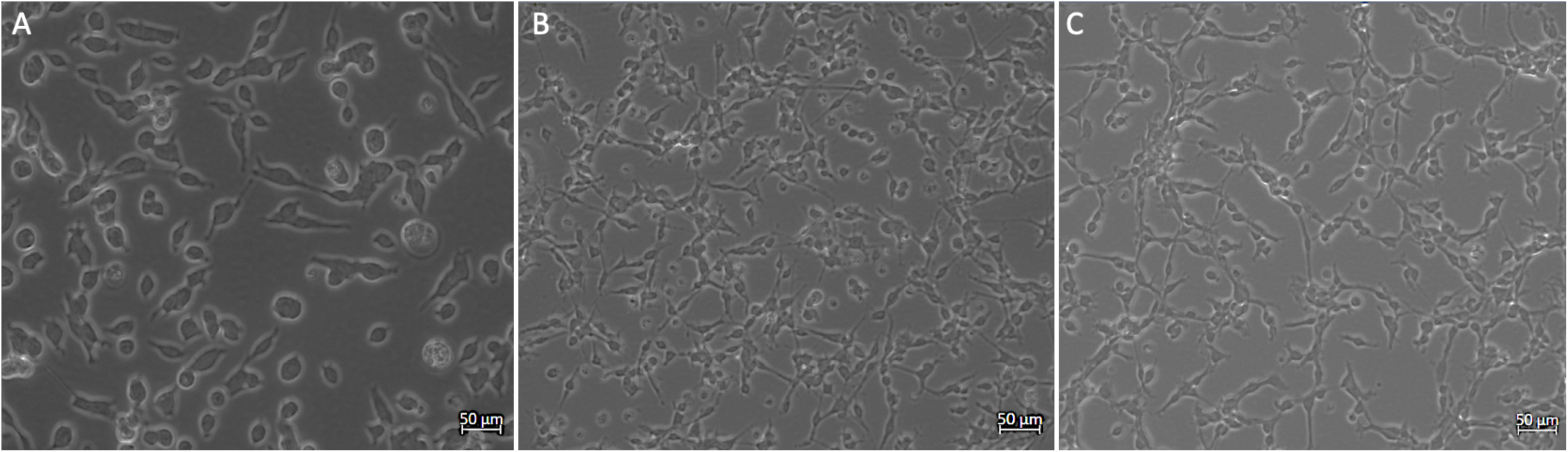
Microscopic analysis of Devil Facial Tumor (DFT) cell lines. Tasmanian Devil Chu-like virus (TDCV) infected DFT1 4906 (A), uninfected DFT1 C5065 (B) and uninfected DFT2 SN (C) cells at confluency. Pictures were taken at 20x after 6 days of growth for DFT1 4906 and 4 days of growth for DFT1 C5065 and DFT2 SN. Microscopic scale is represented at the bottom right of each pictures.

**Figure 3.**
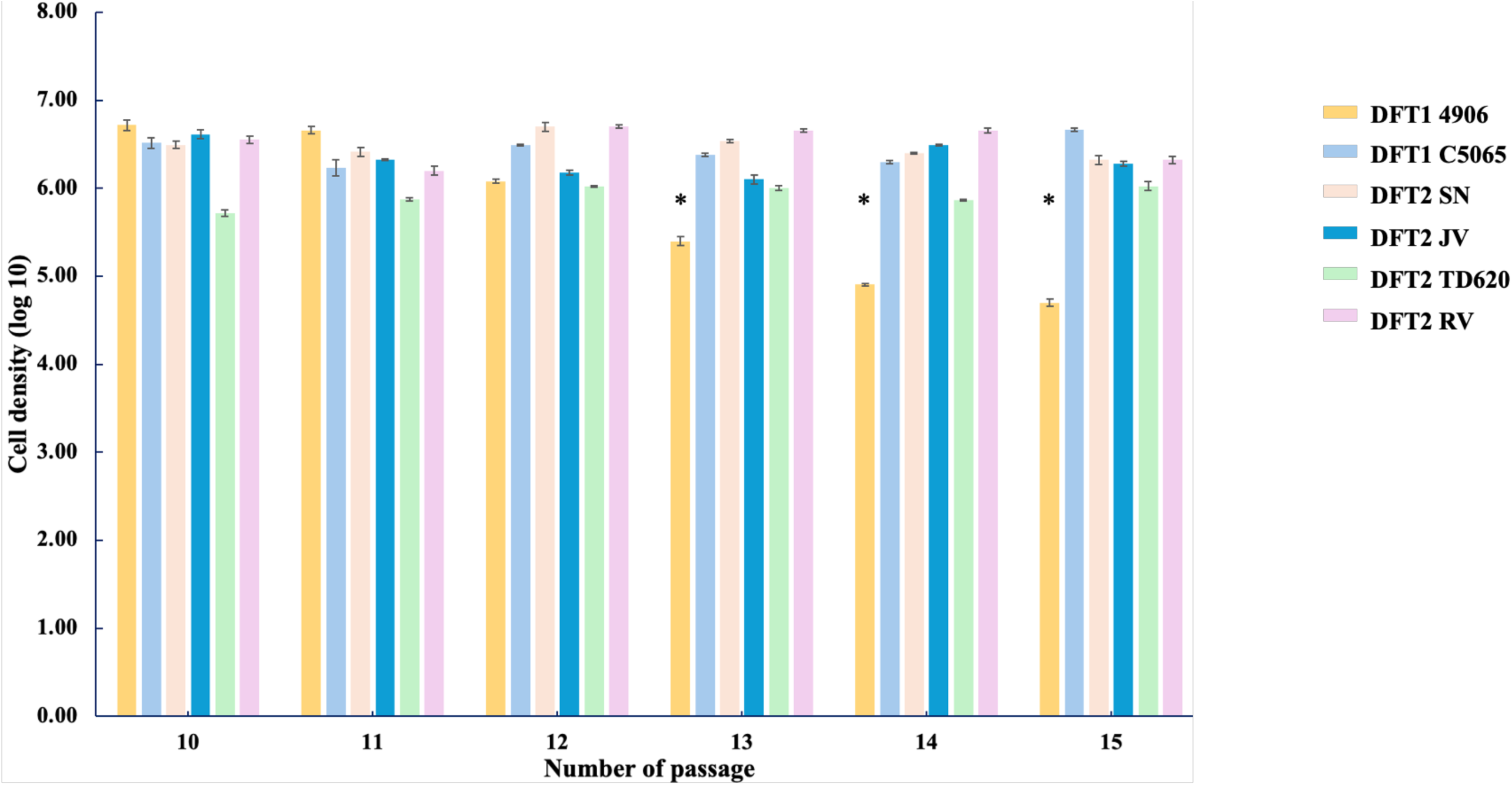
Devil facial Tumor (DFT) cell density in different cell lines after 15 passages. Original aliquot DFT cell lines were passaged 15 times in cell culture. The cell concentration for each cell line was measured after 4-6 days of growth (at confluency) and is expressed as log10 cells/ml (y-axis). Data are represented as mean±SD. Significant differences (*P*-values≤0.05) between cell density in the infected DFT1 4906 cell line and all other non-infected DFT cell lines after several passages are indicated by a * symbol.

### Complete genome sequencing of TDCV from DFT1 4906 cells

To ensure that an infectious, exogenous virus, rather than endogenous chu-like virus elements, were involved with our study, we generated specific primers to amplify and Sanger sequence the full TDCV genome detected in the supernatant of the naturally infected DFT1 4906 cell line. Accordingly, the entire TDCV genome was successfully amplified in three fragments of 4000 bp, 4121 bp and 4167 bp, and Sanger sequencing of the three segments confirmed an intact complete genome of TDCV with the expected, non-disrupted, ORFs. Sequence alignment indicated 100% genomic identity to the original TDCV sequence assembled from the SRA (Library ID SRR6380970) detected from the same DFT1 4906 cell, with no mutations, deletions or insertions.

### Viral replication in Tasmanian devil and mosquito cells

To assess relative viral fitness and exclude that the detection of TDCV was the result of an external contaminate, we first examined the capability of the virus to infect and replicate in new, uninfected tumor cells and fibroblast cells from Tasmanian devils. Accordingly, the tumoral cell line DFT2 SN and fibroblast cell line TD344 were infected with TDCV DFT1 4906 cell culture supernatant for 16 hours, and viral replication was monitored daily. Of note, we observed a significant increase in the viral RNA load of the DFT2 SN cell culture supernatant 6 days post infection (pi) (6.45 log10 copies/mL; + /- 0.1 log10 copies/mL; *p* value <0.05) (**Figure 4A**), but no viral replication in the TD344 fibroblast cells. Interestingly, despite an increase in viral load, no cytopathic effect (CPE) was observed during viral infection of the DFT2 SN cells. The full genome of TDCV was sequenced from the DFT2 SN cell line supernatant six days pi using the same amplification and Sanger sequencing procedures described above. The genome was compared to that observed in DFT1 4906 cell culture supernatant, revealing a total of nine nonsynonymous and two synonymous mutations, of which six were in the polymerase and five in the hypothetical protein (**Figure 5**). To confirm the viral infectivity of TDCV in the new DFT2 SN cell line, the viral supernatant from DFT1 4906 was inactivated by UV and used to infect the DFT2 SN cell line for 16 hours. No viral RNA amplification was observed in the supernatant of the DFT2 cells inoculated by TDCV UV-inactivated 6 days pi and active RNA amplification in non-inactivated supernatant (**Figure 4B**).

**Figure 4.**
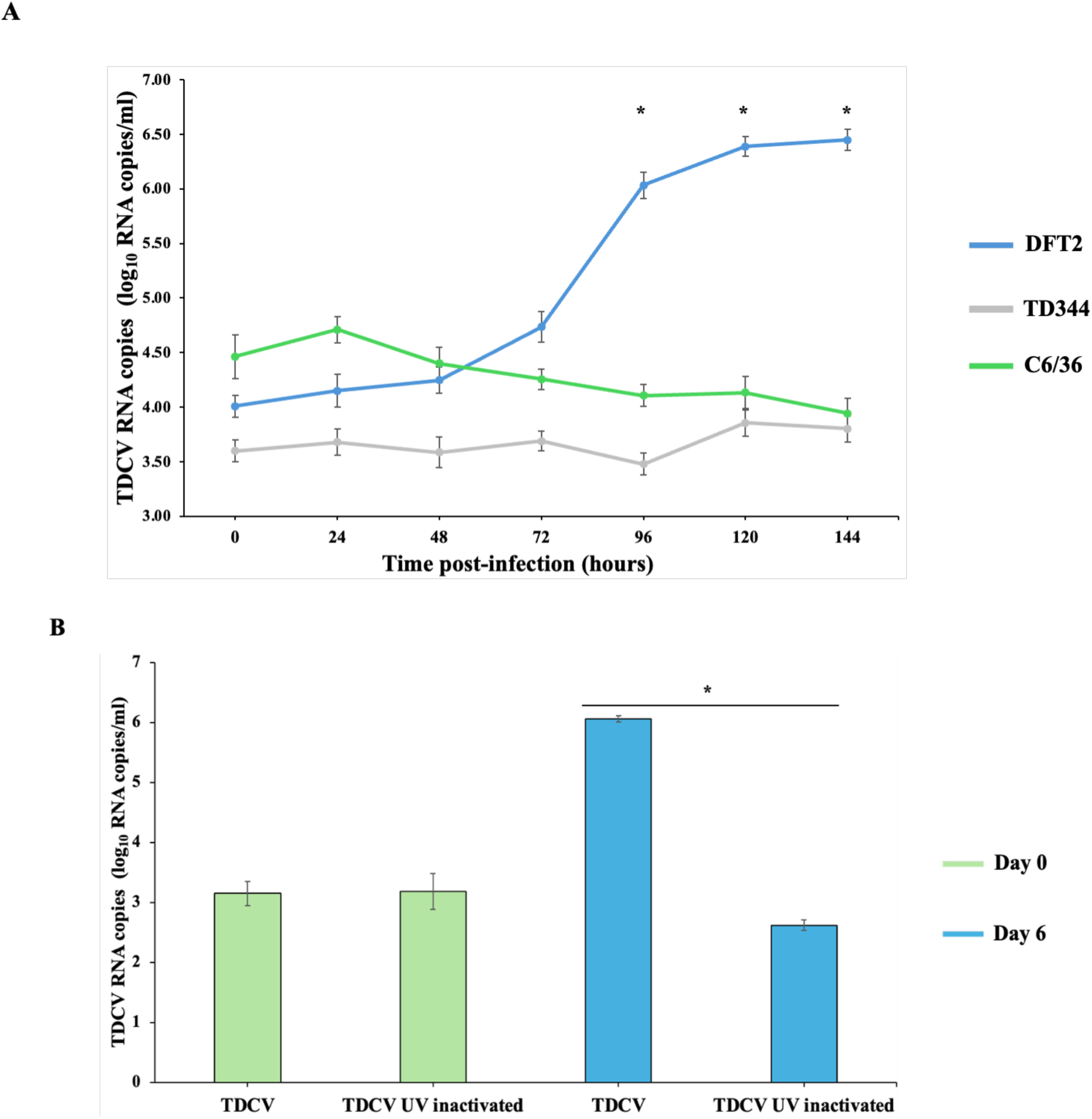
Viral replication kinetics of TDCV in tumor and mosquito cell lines. TDCV replication in additional cell types was assessed by applying a fresh clarified supernatant of TDCV-infected DFT1 4906 cells to DFT2, Tasmanian devil fibroblast (TD344) and mosquito (C6/36) cells (A); or to DTF2 cells with or without UV inactivation prior to application (B). TDCV RNA copies are expressed as log10 RNA copies/ml (y-axis) over time (A) or by treatment (B) (x-axis). Data are represented as mean±SD (indicated by the error bars). Significant differences in viral load (*P*-values≤0.05) are indicated by a * symbol.

**Figure 5.**
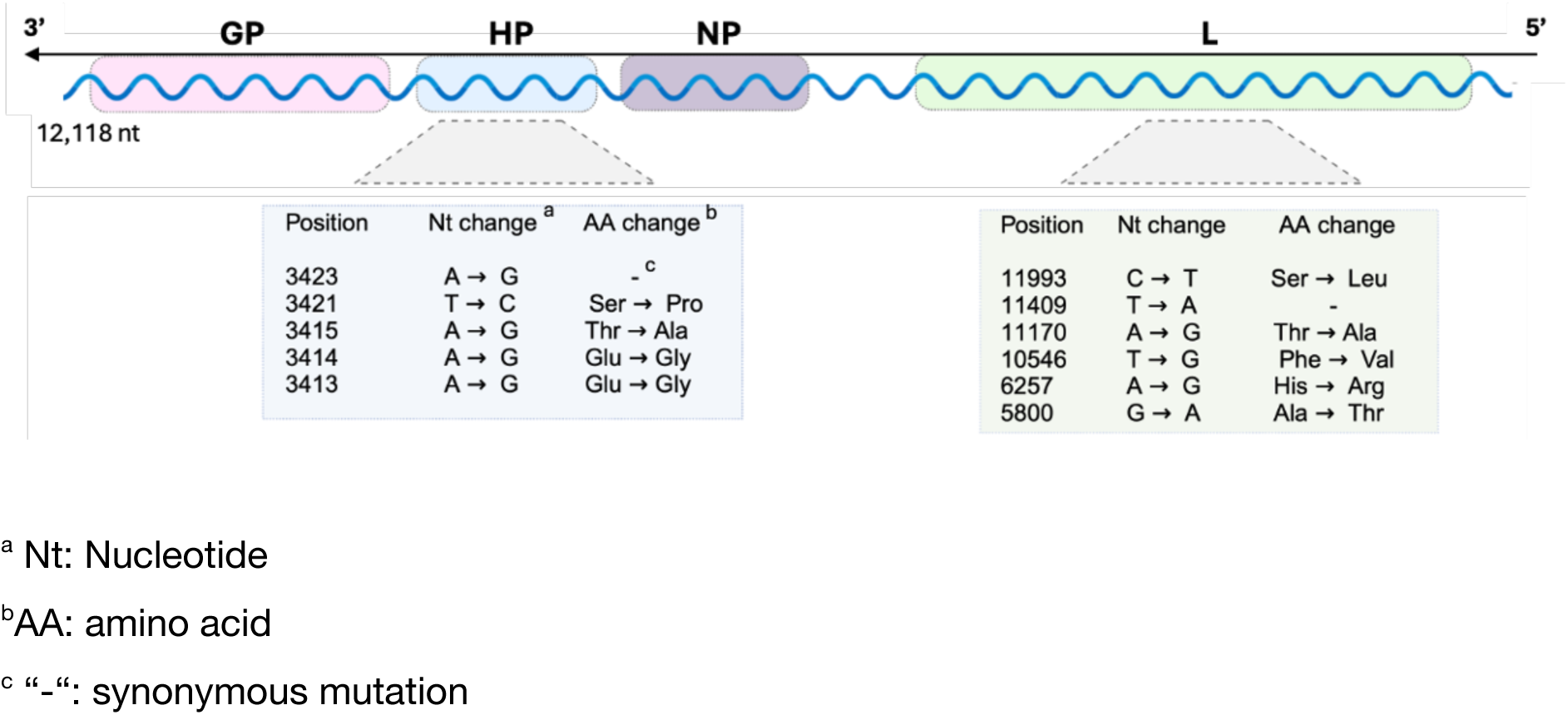
Genome structure of TDCV and mutations observed after passage in DFT2 cells. Genome organization of TDCV in DFT1 4906 cell supernatant. Boxes indicate open reading frames with arrows indicating the direction of translation. GP indicates the putative glycoprotein, NP indicates the putative nucleoprotein and L indicates the putative RdRp. The grey unlabeled box indicates the hypothetical protein of unknown function. Mutations recorded in the TDCV full genome sequence following infection of DFT2 cells (sequenced 6 days post-infection) are listed below the gene in which they occur.

Because chuviruses are widely distributed among arthropods, including mosquitoes, we also analyzed the infectivity of TDCV in the C6/36 *Aedes* mosquito cell line. After six days post-infection we observed no significant changes in the viral RNA load (3.94 log10 copies/mL; + /- 0.14 log10 copies/mL) in cell culture supernatant or cell lysate. Similarly, no CPE was observed on cell monolayers. Taken together, this indicates that TDCV had not undergone replication in mosquito cells (**Figure 4A**).

### Genome annotation

The genome structure of TDCV appeared to be linear and comprise four open reading frames (ORFs) (**Figure 5**). Directly following the 5’UTR region (pos. 1-100 bp) was a putative L-polymerase gene of 6603 bp (pos. 101-6703 bp) followed by the putative nucleoprotein gene of 1260 bp (pos. 7316-8575) and a gene encoding a protein of unknown function of 1071 bp (pos. 8690-9760). A fourth ORF, representing a putative glycoprotein (GP) gene of 2226 bp (pos. 9843-12068), was identified upstream of the 3’UTR region (50 bp; pos. 12069-12118) (**Figure 5**). The putative RdRp and nucleoprotein were detected through comparisons of amino acid sequence similarity with other viruses. Specifically, the conserved region of the RdRp protein had 24% amino acid sequence identity to Lishi spider virus 1 (YP_010839348.1), while across the entire RdRp it shared just 20% amino acid sequence similarity to Hemiperan chu-related virus OKIAV140 (QPL15357.1). The most conserved region of the nucleoprotein had 30% amino acid identity to Salarius guttatus piscichuvirus (UVF58769.1). In contrast, the glycoprotein had no sequence similarity to any reference sequence in NCBI and was putatively assigned because of the length of the ORF, its position in the genome relative to the consensus genome structure of the jingchuviruses, and through sequence alignment to the reference database of jingchuvirus glycoproteins as described below. Interestingly, we also observed a long noncoding sequence of 612 bp just after the putative RdRp (pos. 6704-7315).

### Phylogenetic analysis of TDCV

To determine the evolutionary relationship of TDCV to the broader diversity within the *Jingchuvirales*, we performed a phylogenetic analysis of the putative polymerase, glycoprotein and nucleoprotein amino acid sequences. In all three proteins, TDCV did not cluster with the vertebrate associated piscichuviruses nor with any recently described mammalian associated jingchuviruses (**Figures 6 and 7**). Strikingly, however, in all three proteins it was most closely related (59.9% amino acid identity across the conserved RdRp region and 45% across the entire genome) to a recently identified chu-like virus in RNA-seq data of *Ciona* sp. (sea squirts; i.e. tunicate)^12^ (**Figures 6 and 7**). In the RdRp phylogeny (which is used in species demarcation as per the ICTV guidelines) TDCV and the *Ciona* virus fell between Wufeng rodents chuvirus 1 (unclassified) and a cluster of five sequences including Hubei myriapoda virus 8 (*Myriaviridae*) and Megalopteran chu-related virus 119 (*Crepuscuviridae*) (sequence alignment of 1160 amino acids; **Figure 6, Supplementary** Figure 1). However, this phylogenetic position was unstable and impacted by the length of the sequence alignment (i.e., varied by the stringency of removing poorly aligned regions). A more conservative sequence alignment (336 amino acids) placed TDCV and the *Ciona* virus closer to the cluster of five sequences including the *Myriaviridae* and *Crepuscuviridae* (**Supplementary** Figure 2). Such phylogenetic uncertainty is unsurprising and reflects the long branch lengths within a highly diverse set of viruses. Importantly, however, regardless of alignment length, the mammalian associated viruses do not cluster together and are distributed throughout the phylogeny, distant from the genus *Piscichuvirus* sequences, indicative of multiple independent origins (**Figure 6**).

**Figure 6.**
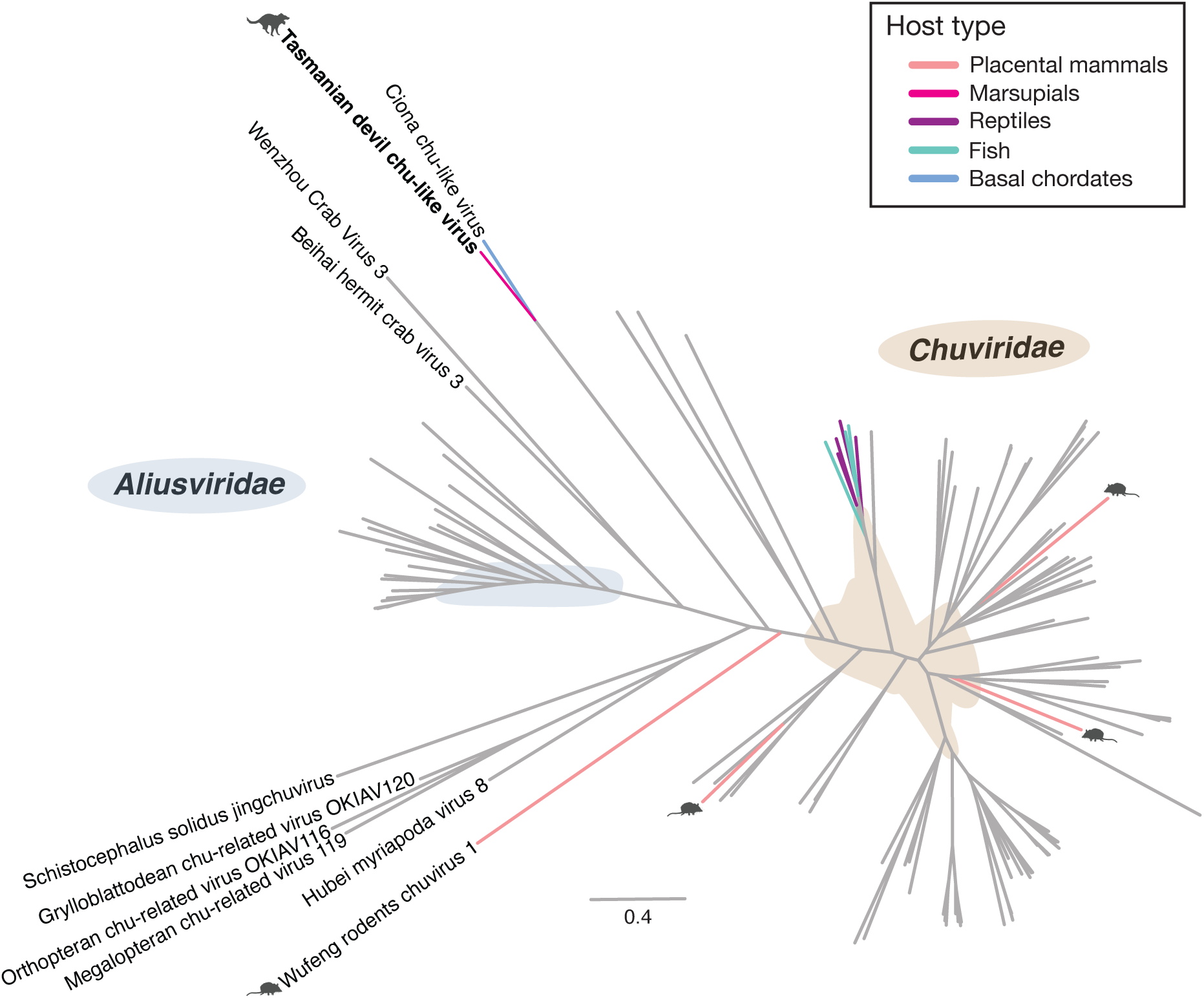
Phylogenetic relationships of the *Jingchuvirales* based on the RdRp. Unrooted maximum likelihood tree of the *Jingchuvirales* RdRp. Chordate associated viruses are indicated with colored tip branches and mammalian associated viruses are indicated with animal silhouettes. The taxon name of TDCV is bolded. Viral families with more than one member are indicated with colored shading and labelled in the figure. The scale bar indicates the number of amino acid substitutions per site.

**Figure 7.**
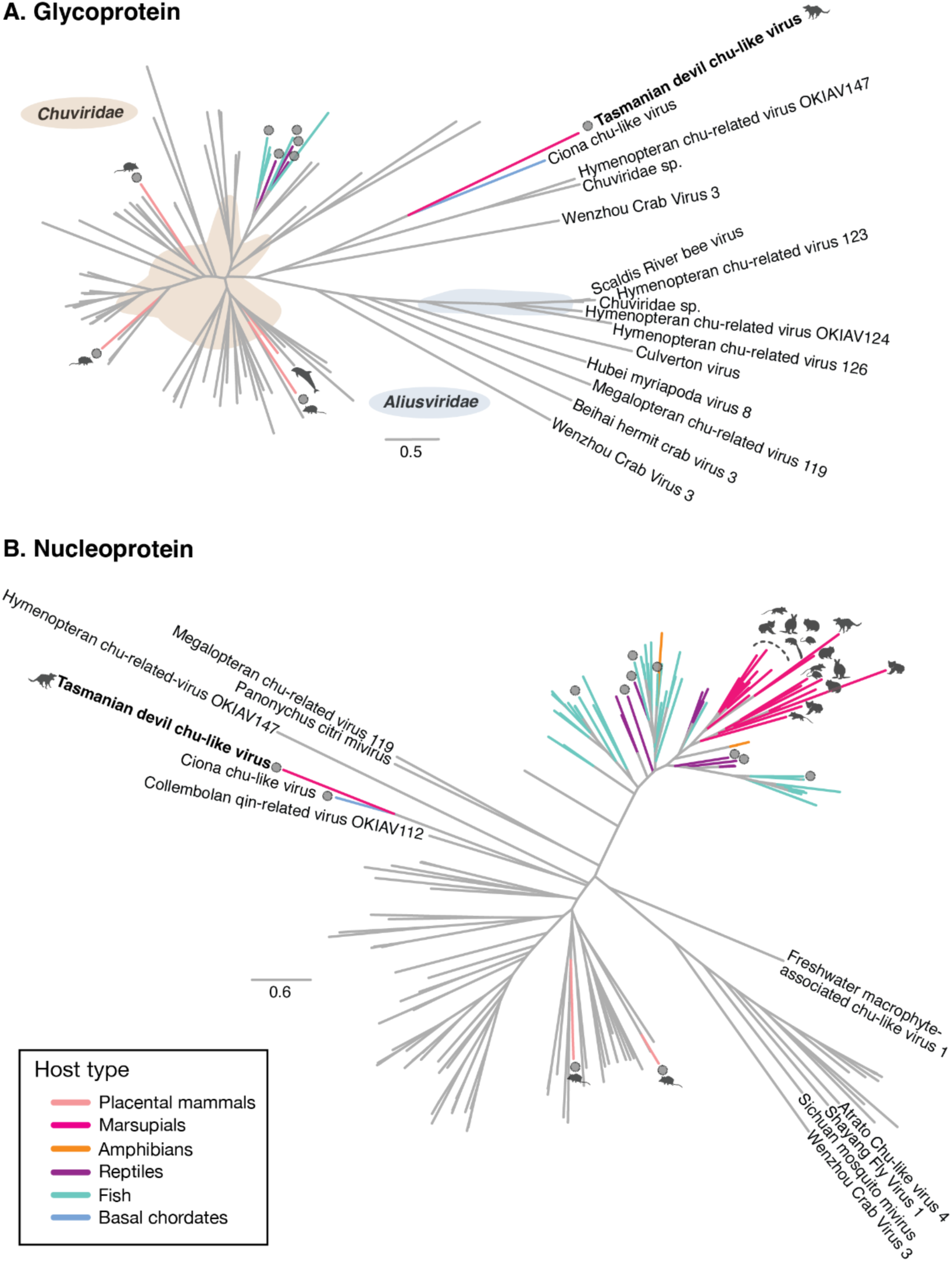
Phylogenetic relationships of the *Jingchuvirales* nucleoprotein and glycoprotein genes. Unrooted maximum likelihood trees of the *Jingchuvirales* structural proteins. Exogenous chordate associated viruses are indicated with a viral particle silhouette at the branch tip and chordate associated virus and EVE sequences are indicated with colored tip branches. Animal silhouettes indicate mammalian associated virus and EVE sequences. The taxon name of TDCV is bolded. Where viral families of more than one member are monophyletic, they are indicated with colored shading and labelled in the figure. (A) Phylogeny of the glycoprotein. (B) Phylogeny of the nucleoprotein (where is no clear clustering of the *Chuviridae* and *Aliusviridae*) . The scale bars indicate the number of amino acid substitutions per site.

TDCV and *Ciona* chu-like virus consistently clustered together in all phylogenetic trees, as did members of the genus *Piscichuvirus*. In the glycoprotein and nucleoprotein phylogenies, TDCV and the *Ciona* chu-like virus were highly divergent from the *Chuviridae* and other defined families (**Figure 7, Supplementary** Figure 3). In addition, because the mammalian associated sequences, which include EVEs, fall in diverse positions, these phylogenies provide further support for multiple host jumping events from invertebrates to mammals (**Figure 7, Supplementary** Figure 4). However, the monophyletic grouping of the piscichuvirus nucleocapsid sequences from EVEs of several marsupial species – including south American marsupials – suggests that an ancient relative of the piscichuviruses infected an ancestor of the extant marsupials. Notably, TDCV does not cluster with these EVEs (**Figure 7**).

## DISCUSSION

We provide experimental confirmation that a novel, highly divergent chu-like virus – TDCV – infects Tasmanian devil (*Sarcophilus harrisii*) cells. In addition, we demonstrate that the host range of exogenous jingchuviruses extends to mammals (marsupials), present the first isolation of a jingchuvirus in cell culture, and the first description of a replicative jingchuvirus. We also demonstrate that not only is TDCV capable of replicating in a tumor cell line (DFT1) where it was originally identified, but that it is also capable of infecting and replicating in a second tumor cell line - DFT2 – that originated in a different Tasmanian devil than, which excludes the probability of virus contamination.

We previously identified TDCV in publicly available RNA-seq data from a Tasmanian devil tumor cell line and hypothesized that the host range of the *Jingchuvirales* extends to the mammals^11^. Subsequent to this publication, four mammalian associated chu-like viruses were identified in rodents and shrews in China through metagenomic sequencing^10^.

However, these viruses were phylogenetically very distant from TDCV, nor did they form a monophyletic group themselves. This supports our hypothesis that the *Jingchuvirales* infect mammals, but suggests that multiple mammalian associated lineages are present within this virus order. The closest relative of TDCV identified to date is in sea squirts (*Ciona* sp.)^12^, implying that there is likely extensive undescribed diversity of *Jingchuvirales* in chordates, which is also supported by the very long branch lengths in all jingchuvirus phylogenies.

Based on the ICTV species demarcation criteria, where a novel species is defined as having <90% amino acid similarity in the L protein (that contains the RdRp), TDCV represents a novel species distinct from the *Ciona* chu-like virus given that they only share 45% amino acid sequence similarity. Similarly, the clade of TDCV and the *Ciona* chu-like virus share only 20% amino acid sequence similarity with their closest relatives in the L protein, meeting the ICTV demarcation criteria (<21% amino acid similarity) for a new virus family. Hence, we tentatively name this family *Lutruwitaviridae* in recognition of the traditional name of the island on which the virus was first identified.

The bias in virus research towards anthropogenically significant species, combined with the challenges in identifying highly divergent viruses through sequence similarity alone, has likely limited our ability to identify the true diversity, host range and age of the jingchuviruses. The presence of marsupial EVEs associated with species predating the divergence of Australian marsupials from American marsupials suggests that the ancient exogenous ancestor of this element infected mammals at least 65 million years ago.

However, the limited sampling of marsupials means we cannot exclude that additional exogenous members of this genus that infect marsupials exist. It is also possible that these viruses may have been overlooked in metagenomic virus discovery studies due to the false assumption that they were associated with invertebrates and perhaps of dietary origin. To better understand evolutionary history, further investigations in Tasmanian and Australian wildlife should be performed to identify any viruses that are related to TDCV.

Very little is known about the viral infection of Tasmanian devil tumor cells and the cellular mechanisms involved in these processes, especially during persistent infection which is likely the mechanism here. The difference in morphology between the naturally infected DFT1 4906 cell line, as well as the decreased cell proliferation compared to uninfected cell lines, needs to be further investigated. It is already known that each of the established DFT1 cell lines have various morphologies^20^, and further work should be done to understand whether chronic infection of tumor cells by TDCV has potentially direct or indirect effects on cell biology. It is well known that several viruses, such as the neurotropic Herpes Simplex Virus (HSV-1), can modulate cellular signaling pathways to suppress apoptosis and persistently infect their host without compromising cell viability, but can induce significant changes in the cytoskeleton and altered morphology^21,22^. Although research on DFT1 has shown that tumor cells display many of the characteristics of healthy Schwan’s cells, nothing is currently known about the role of viral receptors among the cells and the cellular machinery that ensures the infection and replication of TDCV.

The association of these viruses with disease and their mechanisms of transmission remain unknown. Rodent and shrew chu-like viruses were identified in liver, lung and spleen, and confirmed through PCR^10^. Of note, no TDCV RNA was detected in the spleen, liver, kidney, colon or lymph nodes of Tasmanian devils. Experimental infection of Tasmanian devil fibroblasts showed a lack of infectivity, suggesting that TDCV may be tumor cell specific, and that TDCV entry occurs via specific cell surface protein and receptor present in the tumor cells, although perhaps not exclusively. An interesting tropism has previously been recorded for aquatic turtle piscichuvirus, in which mRNA was recorded exclusively in the brain of specimens that died of severe meningoencephalomyelitis^9^. Further experiments should be conducted to understand the tropism of TDCV and determine the molecular and cellular mechanisms involved in viral infection using a panel of Tasmanian devil tumor and brain primary tissues, as well as cell lines established from different organs.

Originally established from *Aedes* mosquito larvae, the C6/36 cell line is highly susceptible and recommended for the isolation and propagation of several human and animal arboviruses in cell culture^23^. As shown by the absence of viral RNA load post-infection, TDCV failed to infect and replicate in mosquito cells. While this result suggests a degree of host specificity, non-insect cell lines such as Vero, and cell lines from other arthropods such as ticks^24^, should be used in the future.

As TDCV falls in a divergent phylogenetic position it is impossible to determine if this virus came directly from an invertebrate, had an invertebrate vector, or came from a currently unsampled vertebrate lineage. To date, there is little evidence on whether and how chuviruses are transmitted from one individual to another. In addition, it is unknown whether viral transmission occurred in parallel to the transmission of DFTD between animals^25^. Of note, ectoparasites are commonly observed on wild devils^26,27^. Most recently, a devil with DFT1 was found to be severely co-infected with both *Sarcoptes scabeii* and *Demodex* mites^28^. Future studies should collect invertebrates during routine Tasmanian devil trapping to facilitate TDCV, which may provide additional context to understand the evolutionary origins and transmission mechanisms of this virus. Combined PCR and serological surveillance will also provide key data on whether TDCV infection or co-infections are associated with DFTD pathology and how the infection modulates the pathology and host cellular and molecular response.

In sum, the combination of experimental and phylogenetic results obtained here, as well the existence of both exogenous and endogenous vertebrate jingchuviruses, strongly suggests that vertebrates, including mammals, are sporadically exposed to jingchuviruses which can occasionally establish sustained transmission cycles. It is perhaps this incipient process of emergence that we have captured with the discovery of TDCV. More broadly, this study paves the way for a better understanding of the highly diverse order *Jinchuvirales* and their host associations. Thanks to the successful isolation of TDCV, experimental studies can now be performed to understand the possible pathophysiology of this Tasmanian devil chuvirus during infection, particularly in the context of DFTD. This work also confirms that an entire order of viruses that were previously only metagenomically described does contain culturable, replicative viruses.

## MATERIALS AND METHODS

### Tasmanian Devil samples

Samples were collected from Tasmanian devils that were euthanized for welfare reasons under animal ethics permit numbers A0012513, A0014976, A0017550 and A26159. The samples were sent from the University of Tasmania to the University of Sydney under the wildlife export permits numbers 2401403 and 2400495 from the Tasmanian government. Samples were collected from 2008 to 2024 across 38 sites in Tasmania (**Supplementary** Figure 5 **and Supplementary Table 1**). Samples were provided by Andrew Flies from the University of Tasmania and Carolyn Hogg from the University of Sydney as either frozen or in RNAlater buffer.

### Cell culture

Six tumor cell lines collected from six different female (DFT1 4906 and C5065) and male (DFT2 SN, JV, RV and TD620) Tasmanian devils (**Supplementary Table 2**) affected by DFTD were cultured at 37°C with 5% CO2 in RPMI 1640 with L-glutamine (1X), 10% heat-inactivated fetal calf serum (FCS; Australian origin), 1% Antibiotic-Antimycotic, 1% HEPES (1M), 1% AmnioMAX™-II, and 50 µM β-mercaptoethanol. Two fibroblast cell lines, the first established from the facial tumor tissue of a Tasmanian devil (named TD344), and a second established from the skin biopsy of the shoulder of another individual (named TD620), were cultured in RPMI 1640 with L-glutamine, 10% heat-inactivated FCS, 1% Antibiotic-Antimycotic, 1% HEPES and 1% AmnioMAX™-II. A mosquito cell line (*Aedes albopictus;* clone C6/36), obtained from the European Collection of Authenticated Cell Cultures (ECACC: 89051705), was cultured at 28°C with 5% CO2 in Eagle’s minimum essential medium (EMEM), 10% heat-inactivated FCS, 2mM Glutamine and 1% Non-Essential Amino Acids (NEAA). Cell viability and density were performed using the trypan blue assay and the percentage of viable cells was calculated with a Countess™ 3 Automated Cell Counter (ThermoFisher Scientific).

### RNA extraction and sequencing

For primary tissues, approximately 10 mg was cut using sterilized scalpels and placed into an RLT lysis buffer (Qiagen) containing 1% β-mercaptoethanol and 0.5% (v/v) Reagent DX (Qiagen). Tissues were then disrupted using the TissueRuptor (Qiagen) for 10 seconds per sample and centrifuged (3 min; 15 000 rpm) to pellet residual tissue, and the supernatant collected. For the cell lines, supernatants were harvested and placed into an RLT lysis buffer containing 1% β-mercaptoethanol and vortex for 1 min. RNA extractions were performed using the RNeasy Plus Mini Kit (Qiagen) following the manufacturer’s recommendation. A DNase treatment was performed to ensure complete elimination of DNA contamination using the QIAGEN RNase-Free DNase Set (QIAGEN) kit following the manufacturer’s instructions. RNA concentrations were then measured using a DS-11 FX+ Spectrometer/Flourometer (DeNovix). Illumina Stranded Total RNA Prep with Ribo-Zero plus (Illumina) was used for preparation of the RNA libraries, and these were subjected to total RNA paired-end sequencing on the Illumina NovaSeq 6000 platform, generating 150 bp reads. Library preparation and total RNA sequencing were performed by the Australian Genome Research Facility (AGRF), Melbourne, Australia.

### Endpoint RT-PCR for TDCV detection in primary tissues and cell lines

We performed RT-PCR to amplify a 1,023 nt fragment of the genome of TDCV using specific primers targeting the putative L gene locus that contains the conserved RdRp. Primer design was based on the TDCV sequence deposited in the SRA database (Library ID SRR6380970), previously detected in the same DFT1 4906 sample from our study. Amplicons were generated using the SuperScript™ IV One-Step RT-PCR System (ThermoFisher) with a mixture containing 12.5 μL of 2X Platinum™ SuperFi™ RT-PCR Master Mix, 1.25 μL of each primer (10 μM), 0.25 μL of SuperScript™ IV RT Mix, 4.75 μL of nuclease-free water and 5 μL of extracted nucleic acids. PCR reactions were performed on a SimpliAmp™ Thermal Cycler at the following conditions: 50°C for 10 min, 98°C for 2 min followed by 35 cycles of 98°C for 10s, 60°C for 10s, 72°C for 30s and 72°C for 5 min. The PCR products were analyzed on SYBR Safe (ThermoFischer) stained agarose gels. Primers are described in **Table 1**.

**Table 1.**
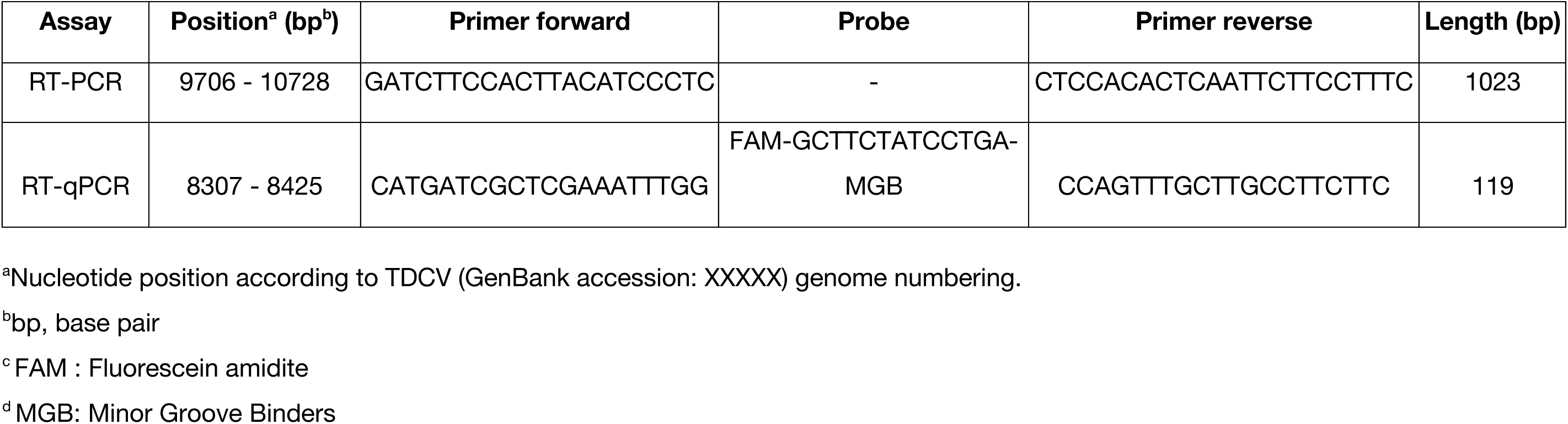
Primers and probes used for TDCV detection by end point RT-PCR and real-time RT-PCR (RT-qPCR). Primers are indicated in the 3’ → 5’ direction of the viral genome.

### Real-time RT-PCR for RNA quantification

The number of TDCV RNA copies was quantified using real time RT-PCR (RT-qPCR). The express GoTaq® RT-qPCR (Promega) was used with 10 μL of GoTaq® qPCR Master Mix, 1 μL of each primer (500 nM), 1 μL of probe (200 nM), 0.4 μL GoScript™ RT Mix for 1-Step RT-qPCR (50X), 1.6 μL of nuclease-free water and 5 μL of extracted nucleic acids in a total volume of 20 μl. Assays were performed using the LightCycler® 480 Instrument II (Roche) with the following conditions: 45°C for 15 min, 95 °C for 2 min, followed by 40 cycles of 95°C for 15s, 60°C for 5s and data collection occurred during the 60°C step. Synthetic RNA was used to calculate the amount of viral RNA from standard curves. Primers and probes used are described in **Table 1**.

### Synthetic RNA transcripts as the gold standard

Synthetic RNA transcript standards were synthesized by *in vitro* transcription using T7 RNA polymerase. First, the template DNA products were produced by RT-PCR from the DFT1 4906 supernatant extracted RNA using a forward primer containing a T7 promoter (**Table 2**). Next, the *in vitro* transcription was performed using the HiScribe™ T7 High Yield RNA Synthesis Kit (NEB). Briefly, 10 μL of the reaction buffer (10X) was mixed with 2 μL of ATP, UTP, GTP, CTP (100mM each), 1 μL of DTT (0.1M), 2 μL of T7 RNA polymerase mix, 2 μL of T7-DNA products (1μg) and 5 μl of nuclease-free water. The reaction was incubated at 37°C for 2 hours and RNA transcripts were then purified using the RNeasy MinElute Cleanup Kit (Qiagen). Concentrations of purified RNA were determined using a DS-11 FX+ Spectrometer/Flourometer (DeNovix).

**Table 2.**
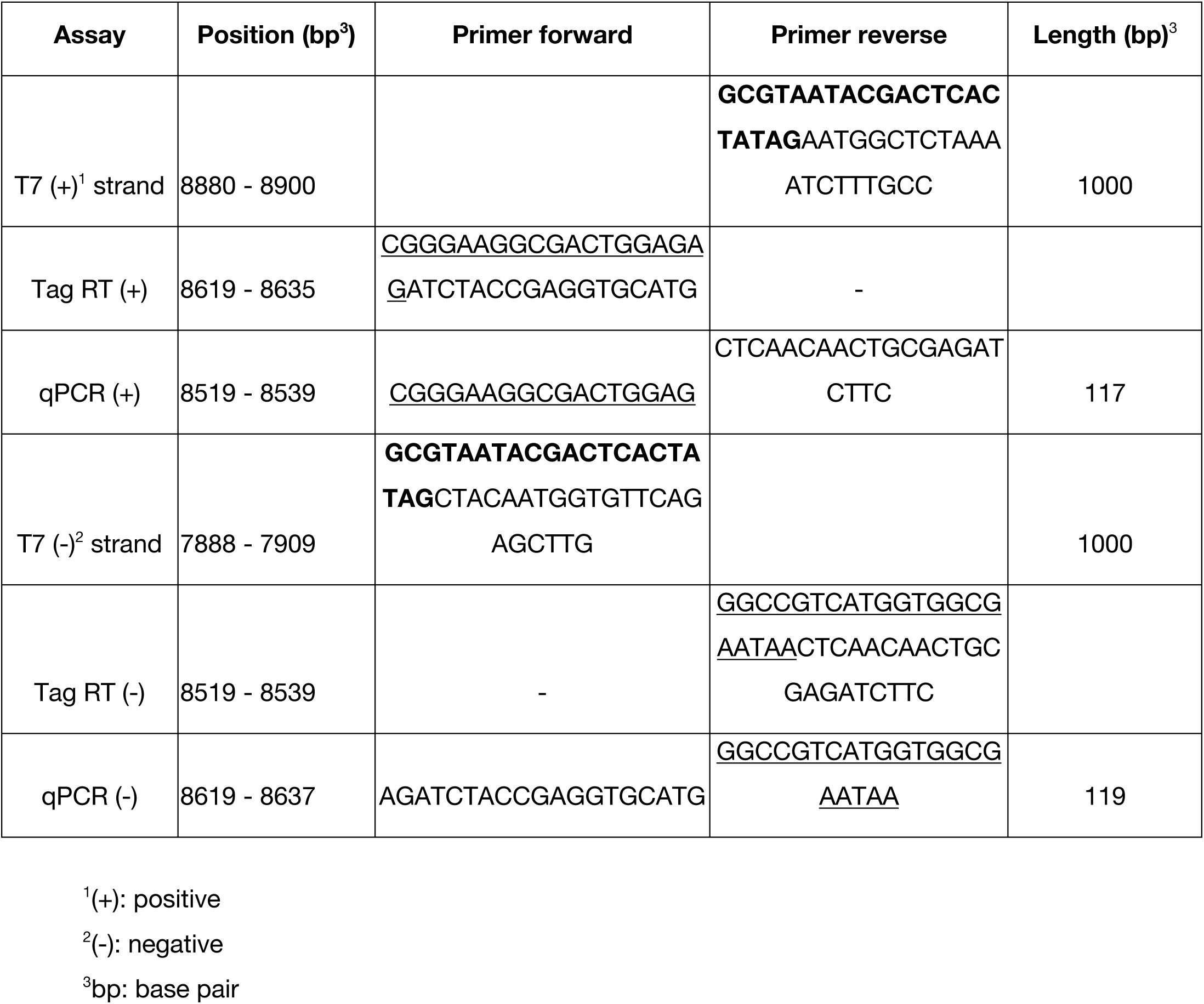
Primers used for the generation of the standard RNA transcripts, RT-PCR tag and strand-specific quantification. Sequences in bold indicate the T7 promoter for the standard RNA generation. Underlined nucleotides represent the unique non-viral tag sequence for the strand specific RT reaction. Primers are indicated in the 3’ → 5’ direction of the viral genome.

### Positive and negative strand labeling and detection

The first positive (+) and negative (-) strand were generated by reverse transcription using the SuperScript™ IV First-Strand Synthesis System (Thermofisher). Reactions were performed using 100 ng of synthetic or viral RNA as template, 1 μL of the appropriate strand-specific primer flanked with a non-viral sequence tag (2 μM) and 1 μL dNTPs (10 mM). The mixture was heated at 65°C for 5 min and incubated on ice for 5 min. In a separate tube, 4μL of SSIV Buffer (5X) was mixed with 1μL of DTT (100mM), 1μL of Ribonuclease Inhibitor and 200 units of Superscript IV Reverse Transcriptase (Invitrogen). The RNA-primer mix, and the RT reaction mix were combined and incubated at 55°C for 10 min and subsequently inactivated by heating at 80°C for 10 min. cDNAs were then used for the endpoint and real-time PCR (qPCR) reaction. The QuantiNova SYBR Green PCR Kit (Qiagen) was used for the strand-specific quantification by real time PCR. The mixture contains 10 μL of SYBR Green PCR Master Mix (2x), 1.25 μL of forward primer and reverse primers (0.7 μM), 6 μL of cDNA and 2.5 μL of nuclease-free water. Assays were performed using the LightCycler® 480 Instrument II (Roche) with the following conditions: 95°C for 2 min, followed by 40 cycles of 95°C for 5s, 60°C for 10 sec, 72°C for 30 sec and data collection occurred during the 60°C step. Synthetic RNA representing each strand, serially diluted from 10^10^ to 10^1^ was used to calculate the number of viral RNA copies. Primers are described in **Table 2**.

### Virus replication kinetics

The infectivity of the TDCV was assessed using (i) a non-infected Tasmanian devil tumoral cell line DFT2, (ii) a Tasmanian devil TD344 fibroblast cell line, and (iii) the mosquito C6/36 cell line. Briefly, a fresh clarified supernatant collected from the second passage of TDCV infected cells (6.5 log10 RNA copies/ml) was diluted in RPMI to 1/10 and a total of 500 μL of the mixture was applied to a 12-well tissue culture plate of DFT2, TD344FBB or C6/36 cells seeded at 5.10^4^ cells/mL one day before. To ensure that the RNA produced is the result of viral replication, an aliquot of the clarified supernatant of TDCV was inactivated by UV-C in a Class II type A2 laminar flow biosafety cabinet (Herasafe; Thermofisher) for 30 min before being diluted and applied to a separate 12-well tissue culture plate of DFT2 cells. Infections were left overnight at 37°C, and inoculant was removed. Cells were then washed 2 times with Hanks’ Balanced Salt Solution (HBSS; ThermoFisher) and 1 mL of culture medium with reduced FCS (2%) was added to the cell monolayer. Cells were observed microscopically over the course of 6 days and one single well from each 12-well tissue culture plate was harvested each 24 h, clarified by centrifugation, aliquoted, stored at - 80°C.Viral RNA copies/mL were quantified using a real-time system.

### RNA-Seq analysis

Sequence reads were assessed for quality using Fastqc, trimmed to remove adapters and low-quality reads using Trimmomatic (v. 0.39)^29^, reads were then assembled *de novo* using Megahit (v.1.2.9)^30^. Assembled contigs were screened for sequence similarity to viruses using a custom virus discovery pipeline as described previously^11^. Briefly, contigs were searched against a custom virus sequence database using Diamond Blastx (v.2.1.6)^31^, contigs with sequence similarity were extracted and compared to the NCBI nt (July 2024) database using Blastn (Blast+ v. 2.12.0)^32^, and all sequences with similarity to non-viral sequences were removed from the data set. The remaining contigs were then compared to the NCBI nr (July 2024) database using Diamond Blastx (v.2.1.6)^31^. Unassembled trimmed reads were also aligned to the TDCV reference genome using Bowtie2 (v.2.2.5)^33^.

### Sanger sequencing

The unpurified PCR products generated by endpoint RT-PCR were directly sequenced by the Sanger method (dual-direction sequencing, AGRF, Melbourne, Australia) using the primers shown in **Table 3**. To correct for any PCR or sequencing errors introduced, the Sanger sequence data generated was then aligned with the Geneious alignment protocol using the default option in Geneious Prime (Geneious Prime® 2025.0.2)^34^. Primer sequences were trimmed from the finalized sequences. The resulting trimmed sequences of the three overlapping amplicons of 4000 bp, 4121 bp and 4167 bp, which together comprise the complete viral genome, were then aligned using the Muscle^35^ method (default options) implemented in Geneious Prime.

**Table 3.**
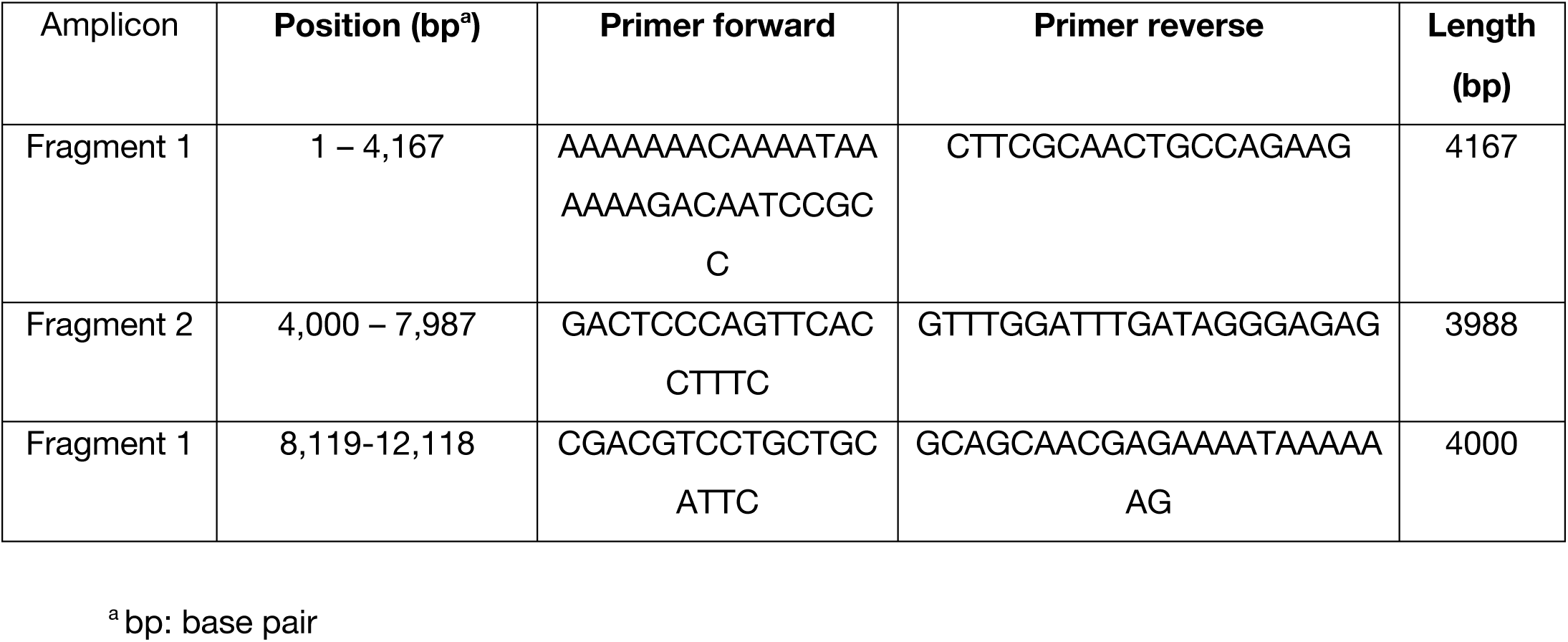
Primers used for end point RT-PCR for amplicon sequencing of TDCV in three overlapping DNA fragments. Primers are indicated in the 3’ → 5’ direction of the viral genome.

### Genome annotation and phylogenetic analysis

The full-length genome sequence of TDCV amplified from the infected cell line (DFT1 4906, CVCL_LB78) was assembled as described above and compared to the genome sequence identified in our previous study of publicly available RNA-seq data^11^. ORFs were predicted using Geneious Prime^34^ and putative protein functions were identified through a Web Blastp search against the NCBI non-redundant protein database (March 2024). Where a blast search did not return results, the hypothetical protein length and an amino acid sequence alignment was used to identify potential protein function.

A custom reference database of related sequences was constructed using NCBI virus (https://www.ncbi.nlm.nih.gov/labs/virus/vssi/<u>#/</u>; access date 04/07/2024) by retrieving all polymerase, glycoprotein and nucleoprotein sequences classified within the *Jingchuvirales*. Unclassified sequences related to TDCV were retrieved from NCBI Web Blastx using amino acid sequence similarity. Redundancy was removed using CD-HIT (v 4.6.1)^36^ with a sequence identity threshold of 0.85 and a minimum length of 300. Amino acid alignments were performed using Mafft^37^ (v 7.3) global alignment with 1000 cycles of iterative refinement, and trimmed to removed regions of low-quality alignment using TrimAL (v 1.4.1)^38^ with a gap threshold of 0.8 and similarity threshold of 0.005. For the nucleoprotein and glycoprotein alignments a minimum conserved length of 25% was set when trimming. Two polymerase alignments were produced, one without prescribing a conserved length (i.e., most conservative settings) and a second with 30% of the alignment conserved.

Phylogenetic trees were estimated using the maximum likelihood method in IQ-TREE 2 (v 2.2.2)^39^, with the appropriate amino acid substitution model for each phylogeny selected using ModelFinder^40^: LG+F+I+ρ4 for the polymerase and glycoprotein, and Q.pfam+F+I+ρ4 for the nucleoprotein. Branch support was assessed using 1000 ultrafast bootstrap replicates^41^. Genetic distances were calculated using the Geneious Prime distance matrix function on trimmed alignments.

### Statistical comparisons of cell density and viral growth kinetics

Exploratory analysis was performed using a two-way ANOVA for multiple comparisons with Sidak’s multiple comparisons test. Statistical analysis and graphical representation were performed using GraphPad Prism 7.00. Each experiment was performed in technical triplicates (*N*=3). On the graphs, only *P*-values≤0.05 were indicated by a * symbol. *P*- values>0.05 were considered as not significant and were not displayed.

## RESOURCE AVAILABILITY

### Lead contact

Edward C. Holmes;

### Material availability

The study did not generate new unique reagents or there are no restrictions to availability

### Data and Code Availability

The complete TDCV genome consensus sequence from the cell culture supernatant of DFT1 4906 has been deposited in GenBank under accession number XXXXXX.

## ACKNOWLEDGMENTS

This work was funded by a National Health and Medical Research Council (NHMRC) Investigator grant (GNT2017197) and an Australian Research Council (ARC) Discovery Project grant (DP240101313) to E.C.H., and by the University of Tasmania Advancement Office through funds raised by the Save the Tasmanian Devil Appeal, a Select Foundation Research Fellowship, and The Tall Foundation to A.S.F. We also thank Dr Carolyn Hogg of the University of Sydney for part of the Tasmanian devil primary tissues. We acknowledge the University of Sydney’s high-performance computing cluster Artemis for providing the computing resources used for this study.

## AUTHOR CONTRIBUTIONS

Conceptualization and Methodology, J.M., E.H. and E.C.H. Formal analysis and Investigation, J.M. and E.H. Writing Original Draft, J.M., E.H. and E.C.H. Writing Review & Editing, J.M., E.H., J.E.M., A.S.F. and E.C.H. Resources, J.M.D., A.S.F. and E.C.H. Funding Acquisition and Supervision A.S.F and E.C.H.

## DECLARATION OF INTERESTS

The authors declare no conflicts of interest.

## Notes

### Competing Interest Statement

The authors have declared no competing interest.

