## Supplementary Figures 1-7 for "Isolation of an infectious mammalian chu-like virus from tumor cells of the endangered Tasmanian devil (*Sarcophilus harrisii*)"

### SUPPLEMENTARY FIGURES AND TABLES

**Supplementary Figure 1. Maximum likelihood phylogeny of the *Jingchuvirales* RdRp with 30% of the alignment conserved in TrimAl (1160 amino acids).**
Chordate associated viruses are indicated with colored tip branches and TDCV is indicated by the Tasmanian devil silhouette. The scale bar indicates the number of amino acid substitutions per site and the tree is midpoint rooted for clarity. Ultrafast bootstrap values >50% are indicated with circles at the node.

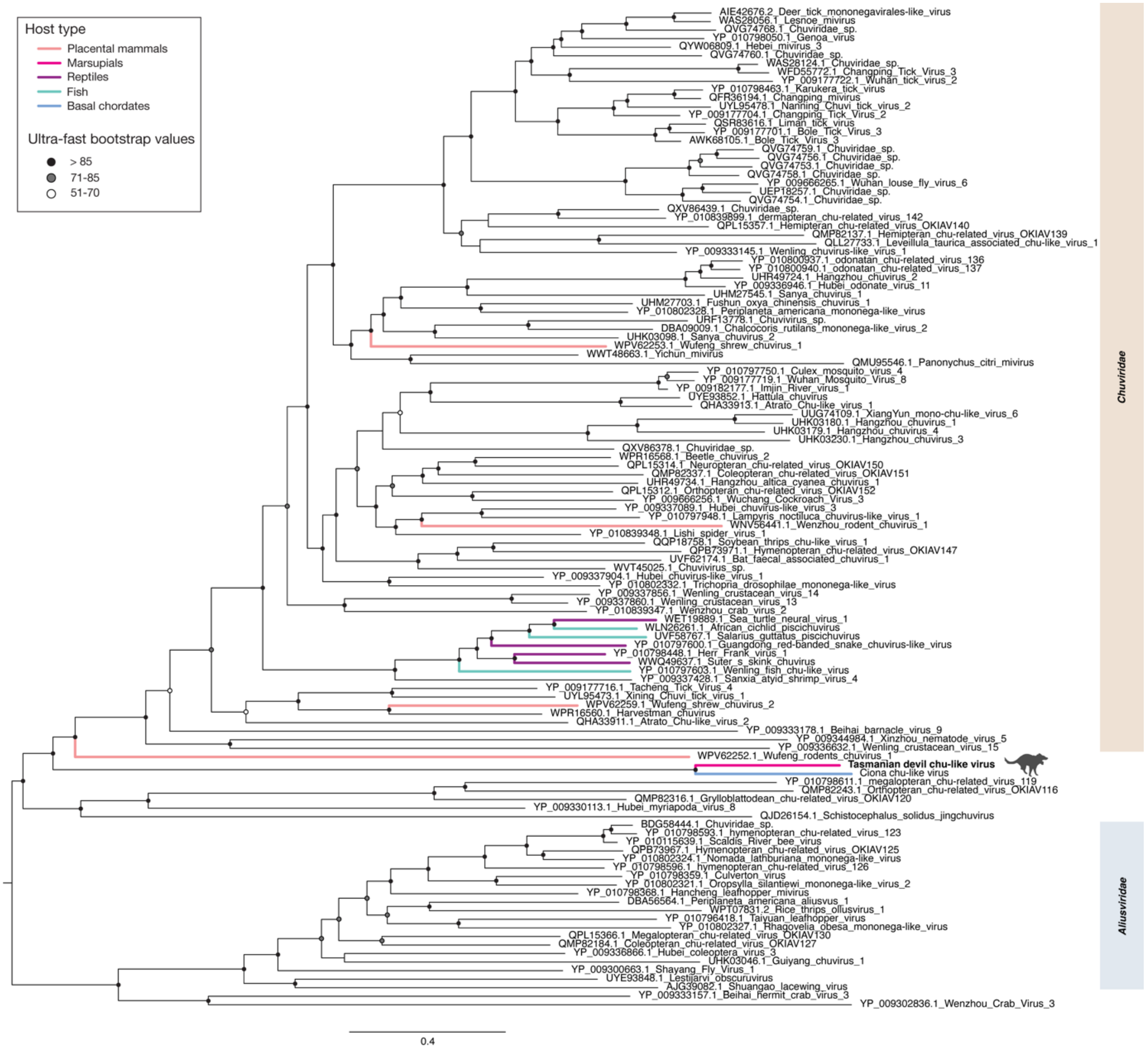

**Supplementary Figure 2. Maximum likelihood phylogeny of the *Jingchuvirales* RdRp with no forced conservation of alignment length in TrimAL (336 amino acids, most conservative setting).**

Chordate associated viruses are indicated with colored tip branches and TDCV is indicated by the Tasmanian devil silhouette. The scale bar indicates the number of amino acid substitutions per site and the tree is midpoint rooted for clarity. Ultrafast bootstrap values >50% are indicated with circles at the node.

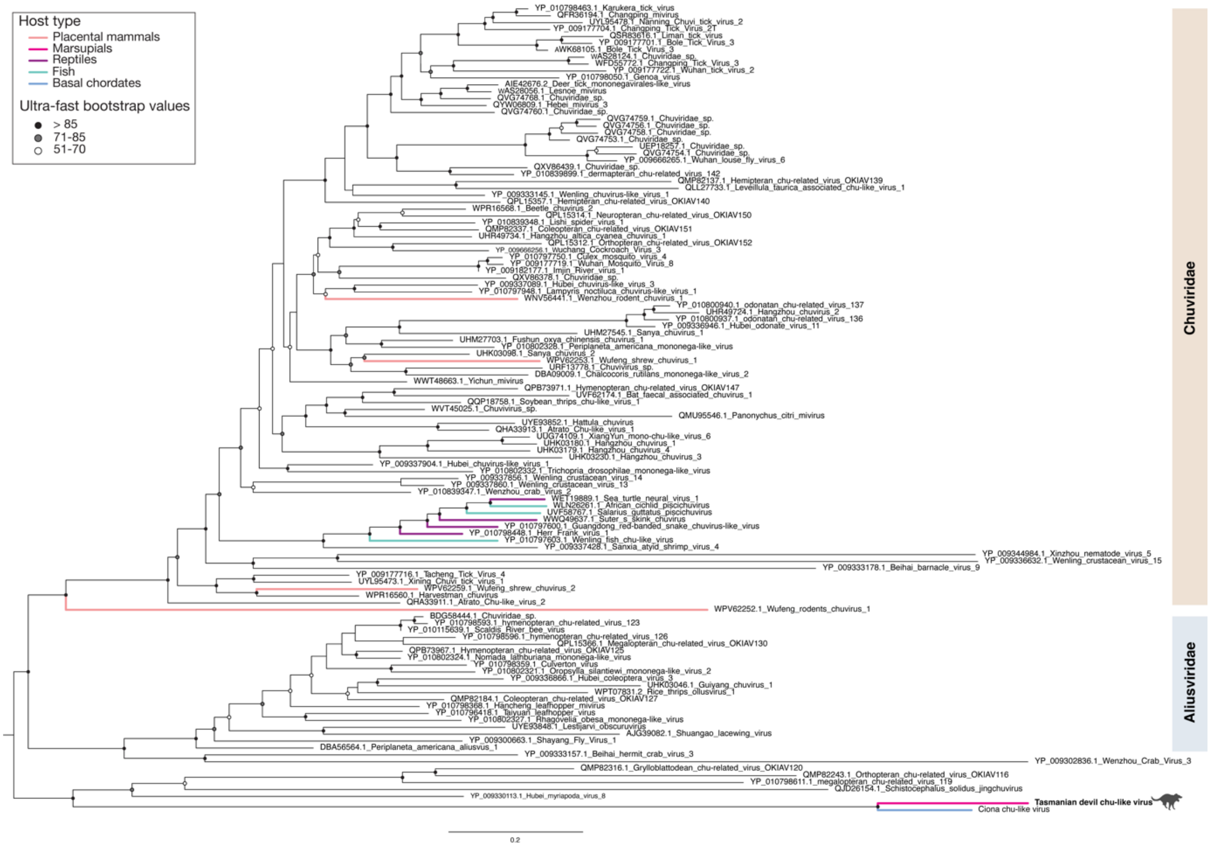

**Supplementary Figure 3. Maximum likelihood phylogeny of *Jingchuvirales* glycoprotein with 25% of the alignment conserved in TrimAl (528 amino acids).** The scale bar indicates the number of amino acid substitutions per site and the tree is midpoint rooted for clarity. Ultrafast bootstrap values >50% are indicated with circles at the node.

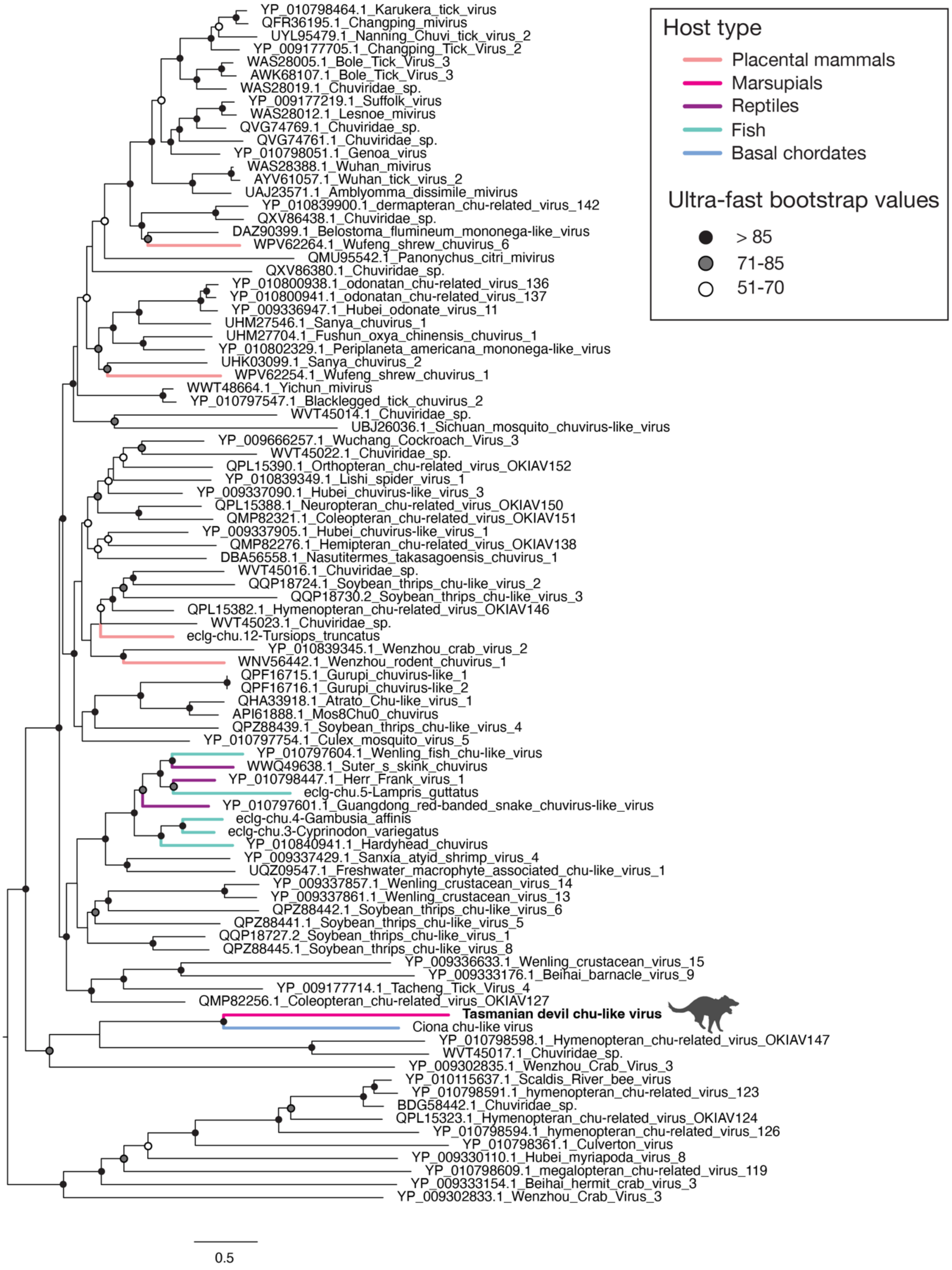

**Supplementary Figure 4. Maximum likelihood phylogeny of *Jingchuvirales* nucleoprotein with 25% of the alignment conserved in TrimAl (594 amino acids).** The scale bar indicates the number of amino acid substitutions per site and the tree is midpoint rooted for clarity. Ultrafast bootstrap values >50% are indicated with circles at the node.

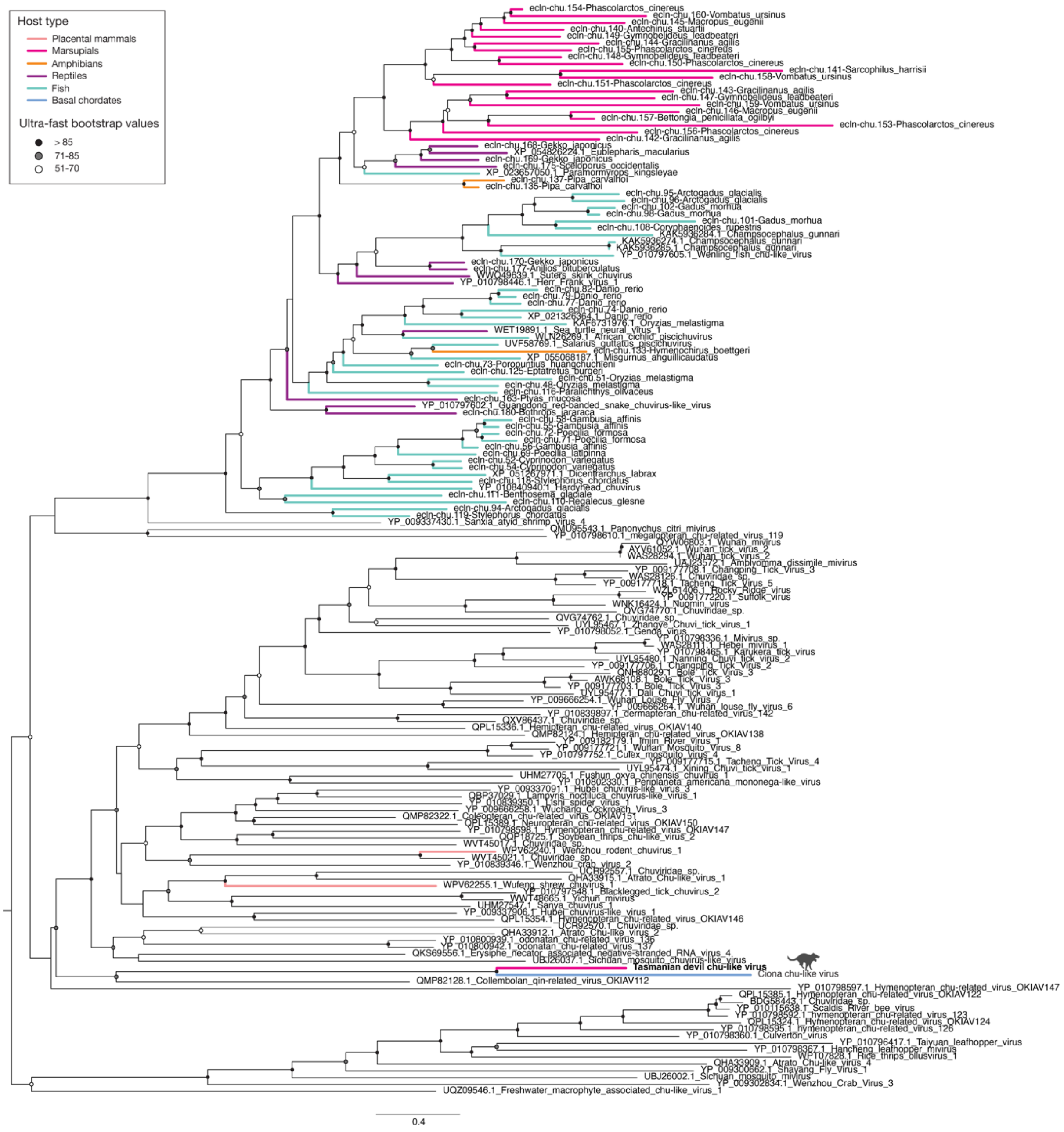

**Supplementary Figure 5. Tasmanian devil tissue sampling sites in Tasmania.**

The corresponding tissue to each site and additional details of each tissue is listed in Supplementary Table 1. The sampling location of the TDCV positive tumor cell line is indicated with a schematic image of cultured cells. Scale of the map is indicated at the bottom left of the picture.

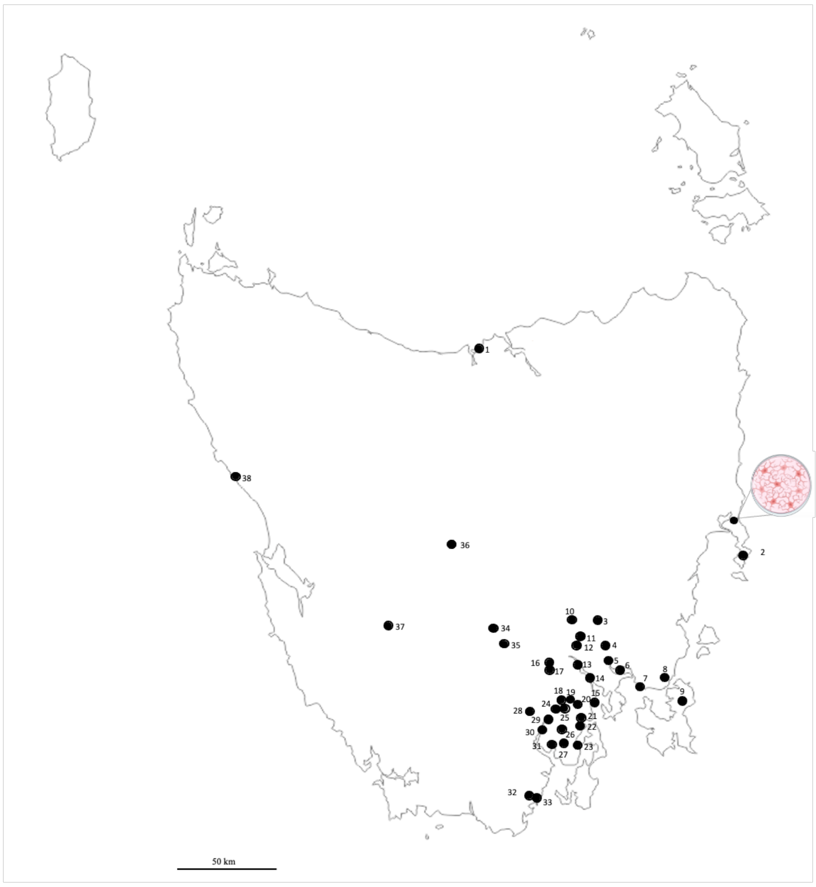


| Site name | Site number |
|---|---|
| Narawantapu National Park | 1 |
| Freycinet Peninsula | 2 |
| Colebrook | 3 |
| Campania | 4 |
| Richmond | 5 |
| Penna | 6 |
| Carlton | 7 |
| Copping | 8 |
| Forestier Peninsula | 9 |
| Kempton | 10 |
| Chauncey Vale | 11 |
| Mangalore | 12 |
| Granton | 13 |
| Risdon | 14 |
| Taroona | 15 |
| New Norfolk | 16 |
| Magra | 17 |
| Mountain River | 18 |
| Leslie Vale | 19 |
| Neika | 20 |
| Margate | 21 |
| Snug | 22 |
| Woodbridge | 23 |
| Grove | 24 |
| Lower Longley | 25 |
| Snug Tiers | 26 |
| Nichols Rivulet | 27 |
| Judbury | 28 |
| Huonville | 29 |
| Franklin | 30 |
| Cygnet | 31 |
| Hastings | 32 |
| Southport | 33 |
| Ellendale | 34 |
| Fentonbury | 35 |
| Bronte Park | 36 |
| Denison river | 37 |
| Granville Harbour | 38 |

**Supplementary Table 1. Details of Tasmanian devil primary tissues sampled across multiple sites in Tasmania from 2008 to 2024 for TDCV and chu-like virus detection.**

| Site | Colon | Kidney | Liver | Lymph nodes | Spleen | Tumor | Total |
|---|---|---|---|---|---|---|---|
| Bronte Park | | | 2 | | 2 | 8 | 12 |
| Campania | 1 | | 1 | | 2 | | 4 |
| Carlton | | | | | 1 | 1 | 2 |
| Chauncey Vale | | | | | 1 | | 1 |
| Colebrook | | 2 | 2 | | 2 | | 6 |
| Copping | | 1 | 1 | | 1 | | 3 |
| Cygnet | | | 1 | | 1 | 1 | 3 |
| Denison river | | | 1 | | | 1 | 2 |
| Ellendale | | | | 1 | | | 1 |
| Fentonbury | | | | | | 1 | 1 |
| Forestier Peninsula | | 4 | 3 | | | | 7 |
| Franklin | | | | | 1 | | 1 |
| Freycinet Peninsula | | | | | | 1 | 1 |
| Granton | | 1 | 1 | | 1 | | 3 |
| Granville Harbour | | | | | 1 | | 1 |
| Grove | | 1 | | | | | 1 |
| Hastings | | | | | 1 | | 1 |
| Huonville | | 1 | 1 | | 1 | | 3 |
| Judbury | | | 1 | | | | 1 |
| Kempton | | | 1 | | 1 | 4 | 6 |
| Leslie Vale | | | | | | 1 | 1 |
| Lower Longley | | | | | 1 | | 1 |
| Magra | | 1 | 1 | | | | 2 |
| Mangalore | | | | | 1 | 5 | 6 |
| Margate | | | | | 2 | 2 | 4 |
| Mountain River | | | 1 | | 1 | | 2 |
| Narawntapu | | | | | | 2 | 2 |
| Neika | | | | | 1 | 4 | 5 |
| New Norfolk | | | | | | 1 | 1 |
| Nichols Rivulet | | | | | | 1 | 1 |

| | | | | | | | |
|---|---|---|---|---|---|---|---|
| Penna | | 1 | 1 | | 1 | | 3 |
| Richmond | | | | | 2 | 1 | 2 |
| Risdon | | | | | 1 | 1 | 2 |
| Snug | | 1 | | | | | 1 |
| Snug Tiers | | | | | 1 | 1 | 2 |
| Southport | | | | | 1 | | 1 |
| Taroona | | | | | | 1 | 1 |
| Woodbridge | | | | | | | 1 |
| Total | 1 | 13 | 18 | 1 | 28 | 37 | 98 |

**Supplementary Table 2. List of Tasmanian devil primary tissues established into tumor cell lines.**

| Site name[a] | Cell type | Name | TDCV RT-PCR[b] |
|---|---|---|---|
| Coles Bay | DFT1 | 4906 | + |
| Bangor | DFT1 | C5065 | - |
| Lower Snug | DFT2 | SN | - |
| Snug Tiers | DFT2 | JV | - |
| Cygnet | DFT2 | RV | - |
| Nicholls Rivulet | DFT2 | TD620 | - |
| Richmond captive | Fibroblast | TD602FBB | - |
| Bronte Park | Fibroblast | TD344FBB | - |

[a] – The name of the site where the animal was captured in order to immortalize the tumor lines
[b] - The cell line that was positive for TDCV by RT-PCR is indicated with a "+" and those cell lines that were negative are indicated with a "-".
